# Low-cost sample preservation methods for high-throughput processing of rumen microbiomes

**DOI:** 10.1101/2022.02.10.480010

**Authors:** Juliana C. C. Budel, Melanie K. Hess, Timothy P. Bilton, Hannah Henry, Ken G. Dodds, Peter H. Janssen, John C. McEwan, Suzanne J. Rowe

## Abstract

**Background:** The use of rumen microbial community (RMC) profiles to predict methane emissions has driven interest in ruminal DNA preservation and extraction protocols that can be processed cheaply while also maintaining or improving DNA quality for RMC profiling. Our standard approach for preserving rumen samples, as defined in the Global Rumen Census (GRC), requires time-consuming pre-processing steps of freeze drying and grinding prior to international transportation and DNA extraction. This impedes researchers unable to access sufficient funding or infrastructure. To circumvent these pre-processing steps, we investigated three methods of preserving rumen samples for subsequent DNA extraction, based on existing lysis buffers Tris-NaCl-EDTA-SDS (TNx2) and guanidine hydrochloride (GHx2), or 100% ethanol.

**Results:** Rumen samples were collected via stomach intubation from 151 sheep at two time-points two weeks apart. Each sample was separated into four subsamples and preserved using the three preservation methods and the GRC method (n = 4×302). DNA was extracted and sequenced using Restriction Enzyme-Reduced Representation Sequencing to generate RMC profiles. Differences in DNA yield, quality and integrity, and sequencing metrics were observed across the methods (p < 0.0001). Ethanol exhibited poorer quality DNA (A260/A230 < 2) and more failed samples compared to the other methods. Samples preserved using the GRC method had smaller relative abundances in gram-negative genera *Anaerovibrio, Bacteroides, Prevotella, Selenomonas*, and *Succiniclasticum*, but larger relative abundances in the majority of 56 additional genera compared to TNx2 and GHx2. However, log_10_ relative abundances across all genera and time-points for TNx2 and GHx2 were on average consistent (R^2^ > 0.99) but slightly more variable compared to the GRC method. Relative abundances were moderately to highly correlated (0.68 ± 0.13) between methods for samples collected within a time-point, which was greater than the average correlation (0.17 ± 0.11) between time-points within a preservation method.

**Conclusions:** The two modified lysis buffers solutions (TNx2 and GHx2) proposed in this study were shown to be viable alternatives to the GRC method for RMC profiling in sheep. Use of these preservative solutions reduces cost and improves throughput associated with processing and sequencing ruminal samples. This development could significantly advance implementation of RMC profiles as a tool for breeding ruminant livestock.

## Background

The microorganisms present in the gastrointestinal tract of ruminants have been widely studied [1, 2] due to their close association with host nutrition [3]. The investigation of this association has been further deepened with the help of genomics and metagenomics [4-6], which has facilitated using information from the ruminal microbiota as a proxy for important characteristics such as feed efficiency [7] and methane emissions [8, 9]. However, the use of ruminal information from metagenomics in breeding for livestock species requires a low-cost and high-throughput approach for generating rumen microbial community (RMC) profiles. Hess et al. [10] investigated the possibility of using Restriction Enzyme-Reduced Representation Sequencing (RE-RRS), also known as genotyping-by-sequencing, to explore the rumen microbial community, and concluded that this genomic tool enabled the investigation of rumen microorganisms (and possibly other environments) with higher throughput and at reduced cost compared to whole metagenome shotgun approaches. RE-RRS is a next generation sequencing technique that reduces genome complexity by digesting genomic DNA using restriction enzymes coupled with sequencing fragments within a certain size range [11]. The success of this technique, however, is dependant on DNA with high quality and integrity.

Preserving rumen samples to maintain DNA integrity for metagenomic sequencing is vital. The standard approach for preserving most microorganisms is by freezing on collection [12]. The Global Rumen Census (GRC) [13] used a protocol developed for frozen samples by Kittelmann et al. [14] to explore rumen microbial profiles of ruminants from across the globe. In this protocol, frozen rumen samples were pre-processed using the two steps of (a) freeze-drying (to preserve and store rumen samples) and (b) grinding (to ensure sample homogenization), before DNA was extracted following the phenol-chloroform with bead beating (PCQI) protocol developed by Rius et al. [15] and modified by Henderson et al. [16]. This rumen sample preservation approach will be termed the “GRC method” throughout our study. A major disadvantage of the GRC method is that the two pre-processing steps of freeze drying and grinding are laborious, costly and low-throughput, hindering application to large-scale studies of RMC profiles.

Studies using targeted sequencing (e.g., from the 16S rRNA gene) show that aspects of rumen samples, such as DNA quality and representativeness of the RMCs, are influenced by time, storage method, sample processing and extraction method [16, 17]. Part of the variation in RMC profiles is related to the particularities of the bacteria that make up the rumen microbiome, which can be classified according to the constitution of their cell wall into gram-negative and gram-positive bacteria. Gram-negative bacteria have a thinner cell wall and are believed to be highly sensitive to freezing and thawing of samples, resulting in degraded DNA [18]. On the other hand, gram-positive bacteria have a cell wall of 10 to 100 times thicker than gram-negative bacteria [19], making it more difficult to disrupt to release DNA.

In this study, we investigated three alternative rumen DNA preservation methods that are based on storing unprocessed rumen samples in a preservative solution. The three solutions we consider are (a) a lysis buffer based on Tris-NaCl-EDTA-SDS (TNx2), (b) a lysis buffer containing guanidine hydrochloride (GuHCl) (GHx2) and 100% ethanol (EtOH). The concentrations for TNx2 and GHx2 were double the final target concentration of the original lysis buffer to ensure an appropriate final concentration after the addition of the rumen sample. TNx2 contains sodium dodecyl sulfate (SDS), a surfactant that has been widely used in molecular biology laboratories but tends to precipitate at room temperature making the process of DNA extraction difficult [20]. The GHx2 solution is similar to TNx2 but uses GuHCl as the salt base instead of NaCl and different compounds are used for the surfactant component. Several studies have investigated the use of these solutions as a preservative using different tissues [21-23]. However, to our knowledge, there are no reports of studies testing these solutions in the concentrations and formulations that we are proposing for investigating ruminal microorganisms. In this study, we investigate the use of these three preservative solutions as alternatives to the GRC method for capturing the RMC profile when using RE-RRS.

## Material and Methods

The use of experimental animals and protocols applied in this experiment were approved by the AgResearch Invermay (Mosgiel, NZ) Animal Ethics committee (approval number 14370). Ewe lambs born in 2017 from the methane selection lines [24, 25] were sampled immediately after measuring methane emissions. Samples were taken at two time-points 14 days apart when animals were approximately 6 months of age. The animals were grazing on ad lib pasture prior to measurements and sampling. Further details about the selection lines, pre sampling grazing management, allocation into groups for sampling and portable accumulation chamber measurements are reported by Jonker et al. [26].

### Solution preparation for rumen DNA storage and lysis

Three solutions were prepared in the laboratory and placed in labelled 8 mL screw cap vials (Sarstedt, Nümbrecht, Germany). The reagents and concentrations of the preservatives were as follows: **TNx2**, 800 mM NaCl, 20 mM Tris HCl, 1.2% of a 10% SDS solution, 200 mM EDTA, pH 8.0 adjusted (with NaOH), 30 ppm ProClin 300 (Sigma-Aldrich, St. Louis, MO, USA); **GHx2**, 2 M GuHCl, 200 mM Tris HCl, 6% Tween20, 1% Triton X-100, 40 mM EDTA, pH 8.0 adjusted (with NaOH), 30 ppm ProClin 300 (Sigma-Aldrich); and **EtOH**, 100% ethanol. Rumen sample fluid is added to the screw capped vials in ratios (sample to preservative solution) of 1:1 for TNx2 and GHx2 and 1:2 for EtOH (Table 1).

**Table 1.**
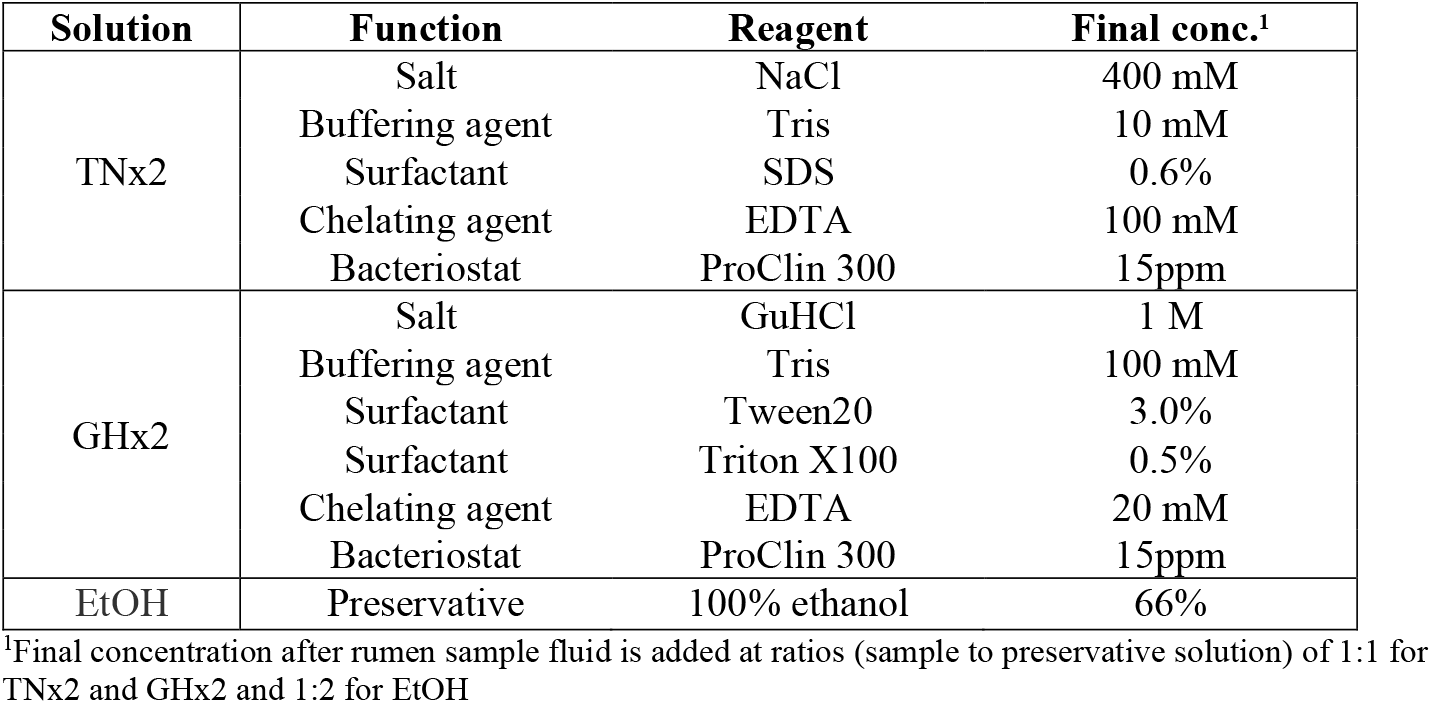
Composition and concentration of preservation solutions after adding rumen sample

### Rumen content sampling

Rumen content (∼25 g) was collected via stomach tubing from 151 ewes, following the standard collection procedure outlined in Kittelmann et al. [14] and placed in an labelled 30 mL pottle (Sarstedt). Three 1 mL subsamples were immediately taken from the rumen contents in the 30 mL pottle using a Gilson pipette with a cut tip (opening bore width ∼5 mm; Thermo Fisher Scientific, Waltham, MA, USA), to ensure a representative sample was taken (fibrous and liquid fraction) and placed into the three 8 mL screw-capped vials containing the TNx2, GHx2 and EtOH solutions. Thus, the rumen sample from a given animal was stored in four separate pottles/vials as follows: 1) ∼25 g of rumen content (without adding solution) placed in a 30 mL pottle (GRC method) and snap-frozen on dry ice; 2) 1 mL of ruminal subsample placed in an 8 mL vial containing 1 mL of TNx2; 3) 1 mL of ruminal subsample placed in an 8 mL vial containing 1 mL of GHx2; and 4) 1 mL of ruminal subsample placed in an 8 mL vial containing 2 mL of EtOH (Fig. 1). The samples in the vials were inverted to mix the contents with the caps screwed on, placed in an insulated container (room temperature or chilled) within 5 minutes of sampling and frozen within 24 hours after sampling. After collection, all samples were stored at −20 °C for approximately 11 months before DNA processing and extraction steps were performed.

**Fig. 1.**
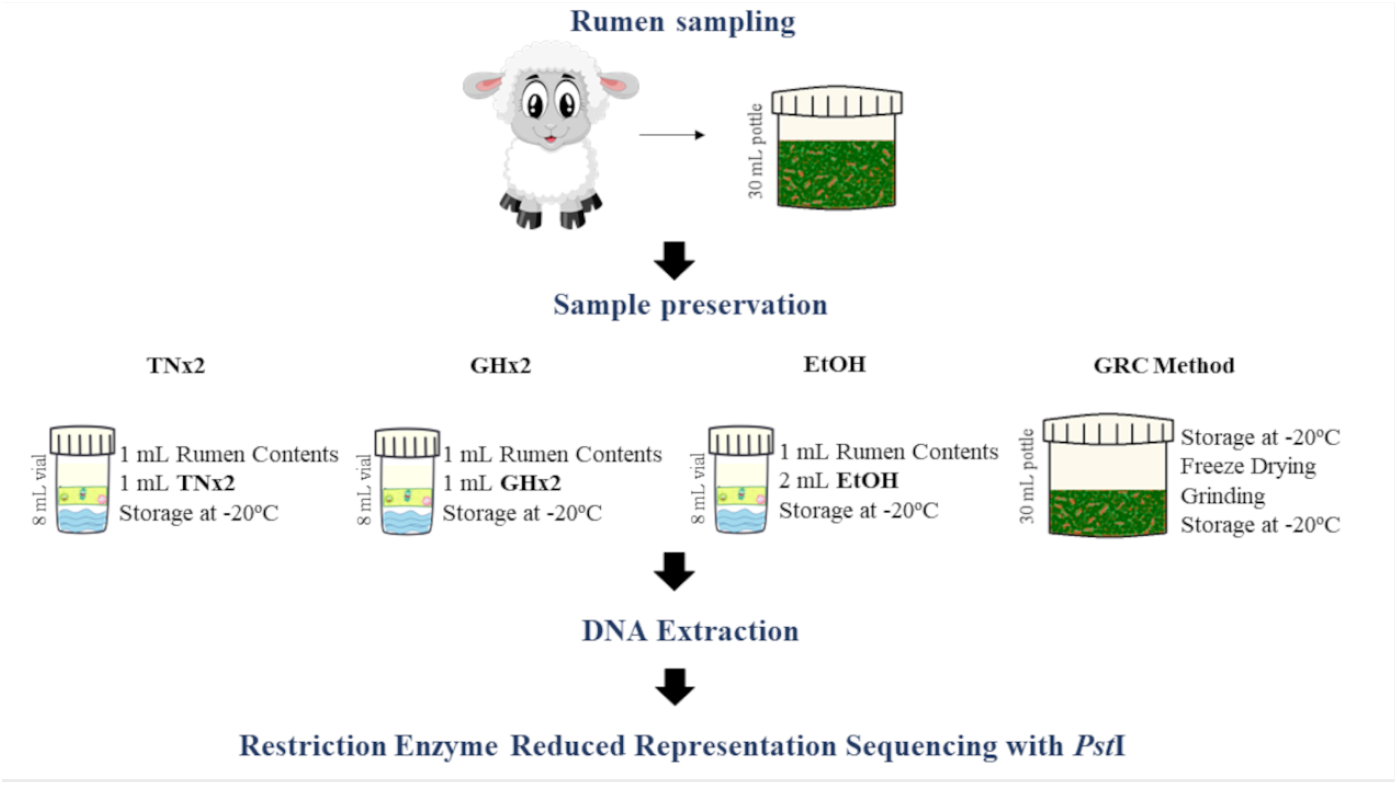
Schematic showing sampling, subsampling and processing of ruminal samples by the four preservation methods

Rumen samples were collected across two days in two rounds approximately two weeks apart (7-8 March and 21-22 March 2018), with each animal sampled once at each round. Thus, 302 rumen samples were taken from 151 animals with each sample divided into 1 pottle and 3 vials, resulting in 1208 rumen subsamples (302 samples of each preservation method) for sequencing and analysis.

### Rumen sample processing

To prepare the 302 samples preserved using the GRC method for DNA extraction, the frozen samples were freeze dried and ground as outlined in the protocol in Kittelmann et al. [14] and Hess et al. [10]. For freeze drying, samples had their lids removed and were placed on stainless steel trays in a −20 °C freezer for approximately four hours then placed in a Gamma 1-16 LSC plus freeze dryer (Christ, Osterode am Harz, Germany) with five shelves [10]. To determine the dry matter (DM) percentage of these samples and to express the DNA yield in µg of DNA per g of ruminal sample dry weight, the samples were weighed before being placed on, and after being removed from, the freeze drier. The percentage of DM was calculated as

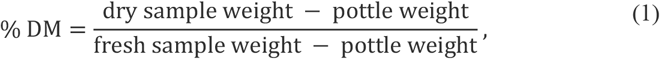

where the pottle weight was set as 8.345 g (the average weight of a 30 mL pottle) for all samples (Additional file 1: Table S1). The freeze drying step plus sample preparation (including weighing) lasted approximately 190 hours, where all the samples were returned to −20 °C after freeze drying. For grinding, batches of samples (∼20 pottles) were removed from the freezer and kept in a cool box with ice packs. Each sample was pulsed for approximately 10 seconds in a Magic Bullet^®^ (NutriBullet New Zealand, Auckland, New Zealand) kitchen blender with cups adapted to handle reduced volumes. At the end of each batch, the samples were returned to -20 °C. In total, it took approximately 32 hours to complete this step, including grinding the samples and cleaning the materials.

For the three alternative solutions, no further sample processing was required. Additional labour consisted of cutting pipette tips for subsampling the rumen content, which took between 5 and 10 minutes for 96 tips.

### PCR plate randomization

Sixteen PCR plates were used for storing DNA extracted from the samples preserved using the four methods. These PCR plates were divided into 4 libraries, each consisting of four plates. Approximately 75 rumen samples preserved using each of the four methods (∼300 samples balanced by preservation method) were randomly assigned into each of the four libraries to form a randomized block design, with an equal number of samples from both sampling rounds present in each library (Additional file 1: Fig. S1, Additional file 2: Data S1). Within a library, individual plates contained samples that were preserved using the same method, while samples corresponding to the same rumen sample (i.e., same animal in the same round) had the same well position across the four plates (e.g., library 1: TNx2 plate: well A1: DNA from animal 875 in round 1; library 1: GHx2 plate: well A1: DNA from animal 875 in round 1; etc.). This randomization was to ensure equity in the results from the four methods tested and to test the library/lane effect during sequencing. Two different positive controls (additional DNA from two GRC samples from round 1) were added to each PCR plate and negative controls were added until all 320 wells of the first 10 columns of the plate were filled (Additional file 1: Fig. S1). DNA extraction for each library was conducted on different days.

### DNA extraction from sheep rumen samples

The exact DNA extraction protocol used was dependent on the method used to preserve the sample (Table 2). For the GRC method, a modified PCQI protocol, first described by Rius et al. [15], was used for DNA extraction. Briefly, DNA extraction from the freeze dried and ground rumen samples was performed through the action of bead-beating in a solution of phenol/chloroform/isoamyl alcohol (25:24:1), a buffer (“buffer A”) composed of NaCl, Tris and EDTA, PM buffer (QIAquick 96 PCR purification kit) and SDS before performing DNA purification using a purification column [15, 16].

**Table 2.** DNA extraction protocols for the four sample preservation methods

| Method | Sample defrosting time | Sample amount used | Rumen sample used (DM) | Solution (μL) |  |  |  | Amount of supernatant taken<sup>5</sup> |
| --- | --- | --- | --- | --- | --- | --- | --- | --- |
|  |  |  |  | Buffer A<sup>1</sup> | Buffer PM<sup>2</sup> | 20% SDS<sup>3</sup> | P:C:I<sup>4</sup> |  |
| GRC | N/A | 30 mg | 30 mg | 282 | 268 | 200 | 550 | ~350 μL |
| TNx2 | 15 min | 1 mL | 34.5 mg | N/A | 268 | N/A | 550 | 500 μL |
| GHx2 | 10 min | 1 mL | 34.5 mg | N/A | 268 | N/A | 550 | 500 μL |
| EtOH | 5 min | 0.8 mL | 18.2 mg | N/A | 268 | 200 | 550 | <350 μL |

<sup>1</sup>200 mM NaCl, 200 mM Tris, 20mM EDTA, pH 8 adjusted with NaOH
<sup>2</sup>Binding buffer (Qiagen)
<sup>3</sup>Sodium dodecyl sulfate (20% wt/vol)
<sup>4</sup>Phenol/chloroform/isoamyl alcohol
<sup>5</sup>Supernatant taken after 4 min of bead beating and centrifugation at 16,060 g and 4 °C for 20 min
N/A denotes not applicable, and DM denotes dry matter

For the preservative solutions, an adaption of the PCQI protocol was used for DNA extraction as follows. First, samples were removed from the freezer and placed on the bench to thaw, where the thaw time varied across the solution types (15 min for TNx2, ∼10 min for GHx2, and ∼5 min for EtOH although the solution was already liquid). After this period, using pre-cut pipette tips as previously described, each sample was vortexed and 0.8 mL (EtOH) or 1 mL (TNx2, GHx2) of the contents of the vial was pipetted and placed into a 2 mL screw-capped vial (Sarstedt) previously prepared with zirconia beads. Physical lysis, as in the GRC method, was performed by the bead-beating technique (Mini-Beadbeater-96, Biospec Products, Bartlesville, OK, USA), and a purification column, using QIAquick 96 PCR purification kit (Qiagen, Hilden, Germany), according to the manufacturer’s recommendations. DNA was eluted in 80 µL of elution buffer (10 mM Tris, pH 8.5 with HCl). As TNx2, GHx2 and EtOH already had preservation and lysis components, only 550 µL of phenol/chloroform/isoamyl alcohol (25:24:1) and 268 µL of the PM buffer were added during bead-beating, while 200 µL of SDS was also added for samples preserved using EtOH (Table 2). After 4 minutes in the “bead-beating” machine and 20 minutes on a Mikro 200 centrifuge (Hettich, Beverly, MA, USA; 16,060 g at 4 °C), between 350 µL and 500 µL of the supernatant were removed for the purification step. One batch of 76 TNx2 samples had to be re-extracted due to errors during DNA extraction. In this case, the remaining samples from the collection pottles were used. DNA extraction for each preservation method was performed separately.

### DNA concentration, purity and integrity

DNA concentrations were quantified by spectrophotometry (NanoDrop TM 8000 Spectrophotometer, Thermo Fisher Scientific), and the apparent specific DNA yield as mg of DNA per g of dry weight of rumen content was calculated considering 6.9% of DM (based on GRC DM; See Additional file 1: Table S1). The relative absorbance readings were investigated for contamination by carbohydrates, aromatic compounds, humic acids and phenol (A260/A230) and proteins (A260/A280) using spectrophotometry. DNA integrity was determined by electrophoresis, on 1% (wt/vol) agarose gel, marked with ethidium bromide, at 100 V and with a 1 kb *Plus* DNA Ladder (Thermo Fisher Scientific) and visualization under ultraviolet light. Twelve samples of each method (48 in total) were checked. Comparative samples (i.e., samples from the same animal and sampling round) were used to load the gel while maintaining the same amount of DNA for the four methods tested.

### Restriction Enzyme-Reduced Representation Sequencing

DNA extracted from the 1208 samples along with the 32 positive controls and 40 negative controls were sequenced using RE-RRS with the restriction enzyme *Pst*I (CTGCA|G) to evaluate RMC profiles across the four methods (GRC, TNx2, GHx2 and EtOH). Hess et al. [10] found this low-cost and high-throughput approach performed similarly to 16S rRNA gene sequencing for capturing the RMC profiles. Briefly, 100 ng of DNA was normalized to 20 ng/µL using PicoGreen (Thermo Fisher Scientific) and digested with the restriction enzyme *Pst*I and ligated with barcodes to link sequences to individual samples [11]. Each library was purified (QIAquick 96 PCR Purification Kit; Qiagen) and the elute PCR amplified using primers and conditions outlined by Elshire et al. [11]. Fragments between 193 and 318 bp were selected by using a Pippin Prep (SAGE Science, Beverly, MA, USA). Each library was checked on a High Sensitivity DNA chip (Agilent Bioanalyzer; Agilent, Satnta Clara, CA, USA), then run on four lanes (one library per lane) of a single flow cell on an Illumina HiSeq 2500 machine (San Diego, CA, USA; High Output Run mode, generating 101 bp single end reads using version 4 chemistry).

### Bioinformatic processing

Sequenced reads were demultiplexed using GBSX [27], with default settings except that mismatches in barcodes or the cut site were disallowed. Reads were then trimmed using cutadapt [28], with a Phred quality score threshold of 20 and a minimum length of 40 bp for individual reads. Samples with fewer than 100,000 reads after demultiplexing and trimming were deemed to have “failed” and were discarded from further analyses. The reference-based (RB) pipeline described in Hess et al. [10], which maps sequenced reads to the bacterial and archaeal genome assemblies of the Hungate1000 Collection [4] with the addition of four *Quinella* genome assemblies [29] and assigns reads to taxonomies at the genus level, was used to generate RMC profiles.

Two RMC profiles were generated: 1) relative abundance matrix, where the number of reads assigned to each genus was divided by the total number of reads assigned to the reference database for that sample, 2) log_10_ relative abundance matrix, in which a pseudo count of one was added to the read counts for each genus before conversion to proportions and then expressed as log_10_ values. Demultiplexed fastq files are available in the NCBI SRA database under BioProject PRJNA791831, and the counts of each genus within each sample used to generate the RMC profile is available in Additional File 2: Data S3.

**Statistical analysis**

Statistical analyses were performed using R v4.0.3 [30]. DNA extraction data were tested by analysis of variance using the R package lme4 v1.1-27 [31] with the statistical model:

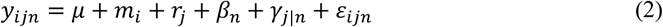

for sample from animal *n* at sampling round *j* and preserved using method *i*, where *y*_*ijn*_ is the DNA yield (spectrophotometry) or DNA quality (A260/A230 or A260/A280), *μ*is the intercept, *m*_*i*_ denotes the preservation method used (GRC, TNx2, GHx2, EtOH), *r*_*j*_ denotes the round the sample was taken, *β*_*n*_ is a random effect for the animal ID such that 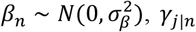 is a nested random effect to account for samples from the same animal taken during the same sampling round such that 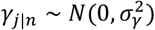, and *ε*_*ijn*_ is random errors such that 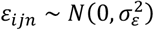. For the RE-RRS metrics (total number of reads and number of reads assigned to the reference database) and log_10_ relative abundances, a fixed effect was included for the library/lane effect (denoted *l*_*k*_) into the statistical model:

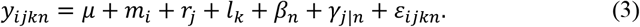

*F*-tests were computed using the R package predictmeans v1.0.6 [32] using a α = 0.05 significance threshold. For the analysis of genus level relative abundances, a Bonferroni correction was applied to the *p*-values using the p.adjust function in R.

A principal component analysis (PCA) was carried out using the prcomp function in R with argument scale=TRUE using the log_10_ relative abundance matrix. Plots of the first three components were colored by preservation method and sampling round. Pearson correlations were computed using the cor function in R for samples collected at the same round but preserved with different methods and for samples collected at different rounds but preserved using the same method.

## Results

### Comparison of methods for DNA quality, DNA quantity and RE-RRS metrics

The apparent specific DNA yield and DNA quality varied depending on the rumen sample preservation and the extraction method used (p < 0.0001) (Table 3, Additional file 1: Table S3). Greater DNA yields were observed for sheep rumen samples preserved and extracted using the GRC method (988 µg/g DM), followed by GHx2 (857 µg/g DM). EtOH performed more poorly on DNA preservation and extraction (419 µg/g DM). DNA quality, although influenced by the preservation method, remained close to the appropriate range for both A260/A230 (2.0 - 2.2) and A260/A280 (1.8 - 1.9) for GRC, TNx2 and GHx2 samples. This suggests low levels of contamination from substances such as carbohydrates (from A260/A230 ratio) and proteins (from A260/A280 ratio). Therefore, the DNA samples were suitable for subsequent analyses, such as PCR (Table 3). EtOH yielded poorer DNA quality compared to the other preservation method with a mean A260/A230 value of 1.96. Sampling round also had significant effects for DNA yield (p = 0.0322) and DNA quality for A260/A230 (p = 0.0004).

**Table 3.** Average DNA yield and quality of sheep rumen samples using four sample preservation methods

|  | GRC | TNx2 | GHx2 | EtOH |
| --- | --- | --- | --- | --- |
| **DNA yield** |  |  |  |  |
| Spectrophotometry<sup>1</sup> | 988 (11)<sup>a</sup> | 554 (11)<sup>c</sup> | 857 (11)<sup>b</sup> | 419 (11)<sup>d</sup> |
| **DNA quality** |  |  |  |  |
| A260/A230 | 2.26 (0.01)<sup>a</sup> | 2.22 (0.01)<sup>b</sup> | 2.25 (0.01)<sup>a</sup> | 1.96 (0.01)<sup>c</sup> |
| A260/A280 | 1.91 (0.01)<sup>a</sup> | 1.87 (0.01)<sup>b</sup> | 1.87 (0.01)<sup>b</sup> | 1.88 (0.01)<sup>b</sup> |
| **Restriction Enzyme-Reduced Representation Sequencing** |  |  |  |  |
| Number of failed samples<sup>2</sup> | 0 | 2 | 3 | 13 |
| Number of reads (thousands)<sup>3</sup> | 1014 (12)<sup>a</sup> | 1004 (11)<sup>a</sup> | 878 (11)<sup>b</sup> | 511 (11)<sup>c</sup> |
| Percent of reads assigned<sup>3,4</sup> | 25 (0.2)<sup>c</sup> | 27 (0.2)<sup>b</sup> | 28 (0.2)<sup>a</sup> | 24 (0.2)<sup>d</sup> |

<sup>1</sup>NanoDrop: µg/g of dry weight rumen content
<sup>2</sup>Sample are consisted failed if there were fewer than 100,000 reads
<sup>3</sup>Excludes failed and control samples
<sup>4</sup>Assigned to the reference database
Values (predicted mean (SED)) in the same row with different letter superscripts denote groups with statistically significant means based on the *F*-test ( $p < 0.05$ ) computed using the predictmeans R package

The DNA integrity, as shown by electrophoresis gel (Fig. 2), revealed that TNx2 and GHx2 were able to yield high molecular weight DNA (above 5 kb which is the recommended level for RE-RRS), which was similar to the results observed with the GRC method (Fig. 2 c). EtOH, however, compromised DNA integrity. Because it was not possible to efficiently preserve and consequently extract DNA from some samples preserved by EtOH, two wells were not able to be loaded with DNA (Fig. 2 b), which further indicates this may be a low-efficiency method compared to the others. Rumen samples preserved with TNx2 and GHx2 had similar color at different steps of the extraction process, while those preserved with EtOH had a darker colour compared to TNx2 and GHx2 (Additional file 3: Figs. S2-S4).

**Fig. 2.**
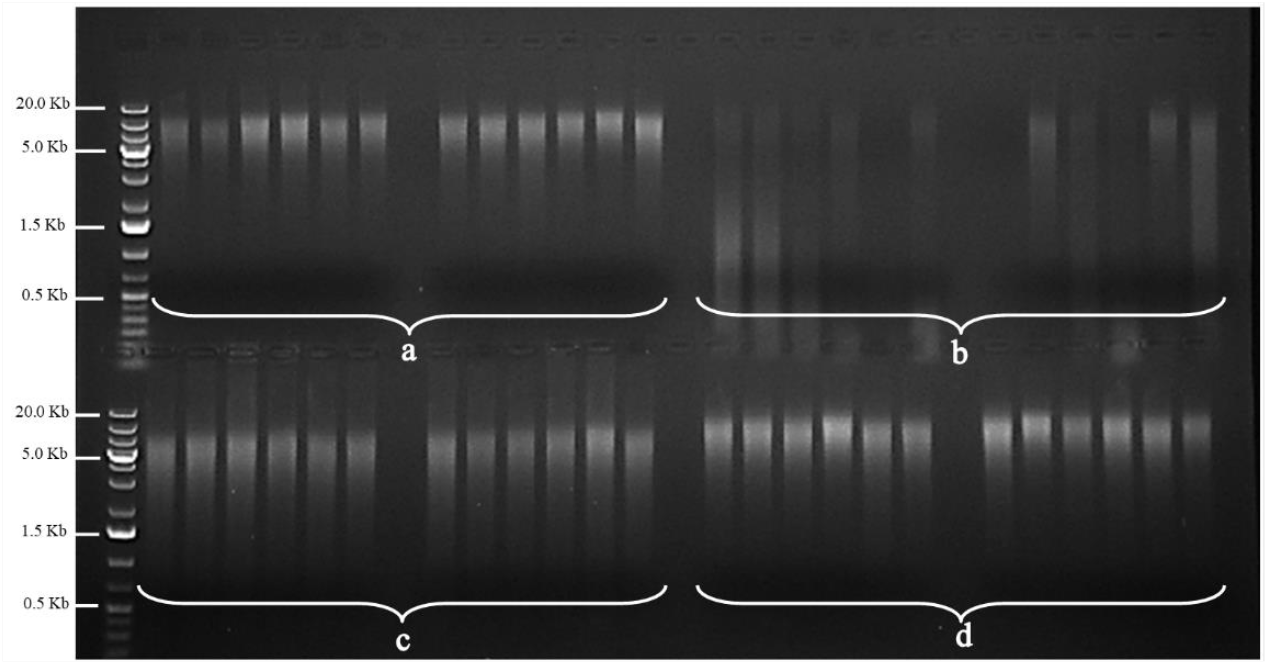
Electrophoresis gel of 12 sheep samples to show DNA integrity from four sample preservation methods. **a** GHx2 **b** EtOH **c** GRC **d** TNx2

**Fig. 3.**
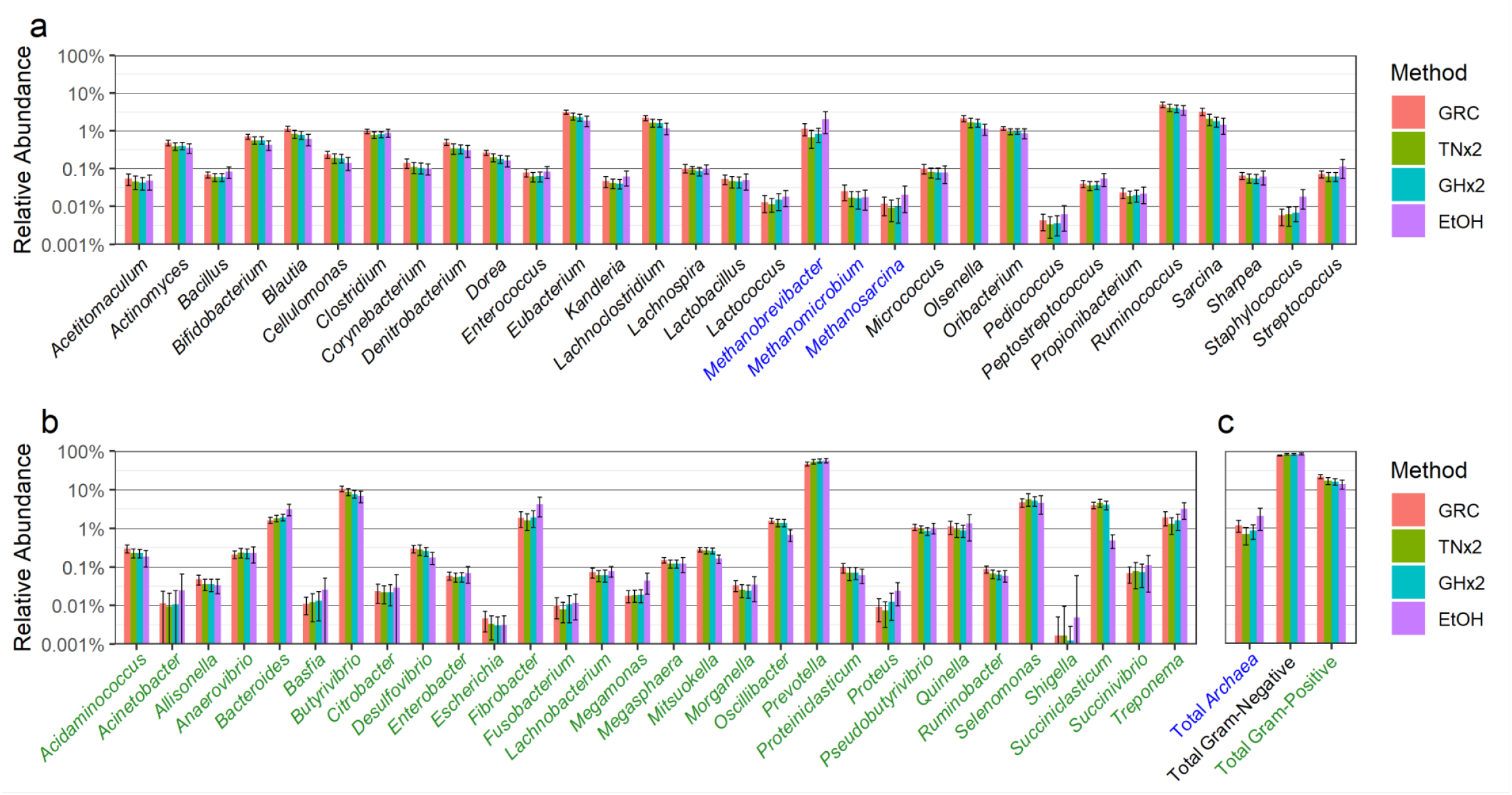
Relative abundance of bacterial and archaeal taxa in samples stored and extracted by the GRC, TNx2, GHx2 and EtOH methods. **a** bacteria with a gram-positive wall (black) and archaea (blue). **b** bacteria with a gram-negative wall (green). **c** Total sum of archaea, gram-positive and gram-negative taxa. Error bars represent ± one standard deviation of the raw means.

For DNA sequencing by RE-RRS, 13 EtOH samples were deemed to have failed (< 100,000 reads), while there were three, two and zero failed samples for GHx2, TNx2 and GRC respectively (Table 3). These failed samples were discarded from further analysis. Preservation method (p < 0.0001), sampling round (p < 0.016) and library/lane (p < 0.001) all impacted the number of reads and reads assigned using the RB approach (Table 3, Additional file 1: Table S3). There were approximately 1 million reads on average for GRC and TNx2 samples, whereas the number of reads for EtOH samples was around half this number (500,000). The percentage of reads assigned to genus level taxonomy using the RB approach was between 24% and 28% across all methods, similar to proportions obtained by Hess et al. [10].

The number of reads for all negative samples was between 50 and 7004, suggesting these samples were true negatives and no significant cross-contamination or reagent contamination had occurred. Positive control samples (across two unique individuals) clustered tightly on a PCA plot (Additional file 3: Fig. S5) and overlapped other samples preserved using the same method (GRC).

### Relative abundance of the bacterial community and archaea

The relative abundances of gram-positive bacteria, archaea and gram-negative bacteria are shown in Fig. 2. The relative abundance of 60 taxa (among the 61 investigated) and the total relative abundance of gram-positive, gram-negative and archaea taxa varied across the four sheep rumen sample DNA preservation methods (p < 0.0001) (Additional file 1: Table S4). *Shigella* was the only taxon that was not significantly different between the preservation methods, largely due to it being the lowest abundant genus observed.

The relative abundances of the 30 gram-positive and the 28 gram-negative bacterial taxa were larger or similar for the GRC method compared to TNx2 and GHx2, except for the five gram-negative bacteria *Anaerovibrio, Bacteroides, Prevotella, Selenomonas* and *Succiniclasticum. Prevotella* was the most abundant taxon (46% - 56%), while *Selenomonas* and *Succiniclasticum* were both moderately abundant (4% - 6%). The relative abundances in these three genera were much lower in DNA extracted using the GRC method. This likely explains the 5% to 6% lower relative abundance of total gram-negative bacteria for the GRC method compared to TNx2 and GHx2. The relative abundance in the three archaea taxa and total archaea were also greater with the GRC method compared to TNx2 and GHx2. Note that here we are using gram-negative in its structural sense, recognising that some genera are phylogenetically gram-positive but have generally gram-negative cell wall structures [33].

The relative abundance of 43 taxa (18 gram-negative, 23 gram-positive and 2 archaea) were also affected by sampling round (Additional file 1: Table S4). Library/lane effect was observed to be significant on the relative abundance of 6 of the 61 taxa (*Cellulomonas, Escherichia, Lachnobacterium, Quinella, Ruminobacter, Succinivibrio)* (Additional file 1: Table S4). Apart from *Quinella* (∼1%), all of these taxa were in low abundance (< 0.12%).

The mean relative abundances for EtOH samples were notably different compared to the GRC, TNx2 and GHx2 samples. In particular, the mean relative abundance decreased considerably for *Oscillibacter* and *Succiniclastium*, and increased notably for *Bacteroides, Fibrobacter*, and *Methanobrevibacter* for EtOH samples compared to the GRC, TNx2 and GHx2 samples. These results, along with the poor DNA yield and quality and lower RE-RRS metrics, suggest that EtOH is less reliable and performs more poorly than TNx2 and GHx2 compared to the GRC method. Hence, EtOH was not considered in any further analyses.

### Mean, variability, and correlation of RMC profiles

Fig. 4 shows pairwise comparisons of the mean and standard deviation of log_10_ relative abundances from the TNx2 and GHx2 preservatives and the GRC method. The mean log_10_ relative abundances were very similar for all three methods, with both the regression coefficients (β) and R^2^ values very close to 1 (Fig. 4 b,c,f). The log_10_ relative abundance standard deviations for TNx2 and GHx2 were larger than for the GRC method (Fig. 4 g,h), as most points were below the red diagonal line of identity. The relationship between the mean and standard deviation of the log_10_ relative abundance was similar across the three methods, with larger variability for lower relative abundances (Fig. 4 a,e,i). This relationship was stronger for the GRC method compared with TNx2 and GHx2, although there were four microbes (*Fibrobacter, Methanobrevibacter, Pseudobutyrivibrio* and *Treponema*) that deviated from the curve with larger standard deviations compared to microbes with similar mean log_10_ relative abundances.

**Fig. 4.**
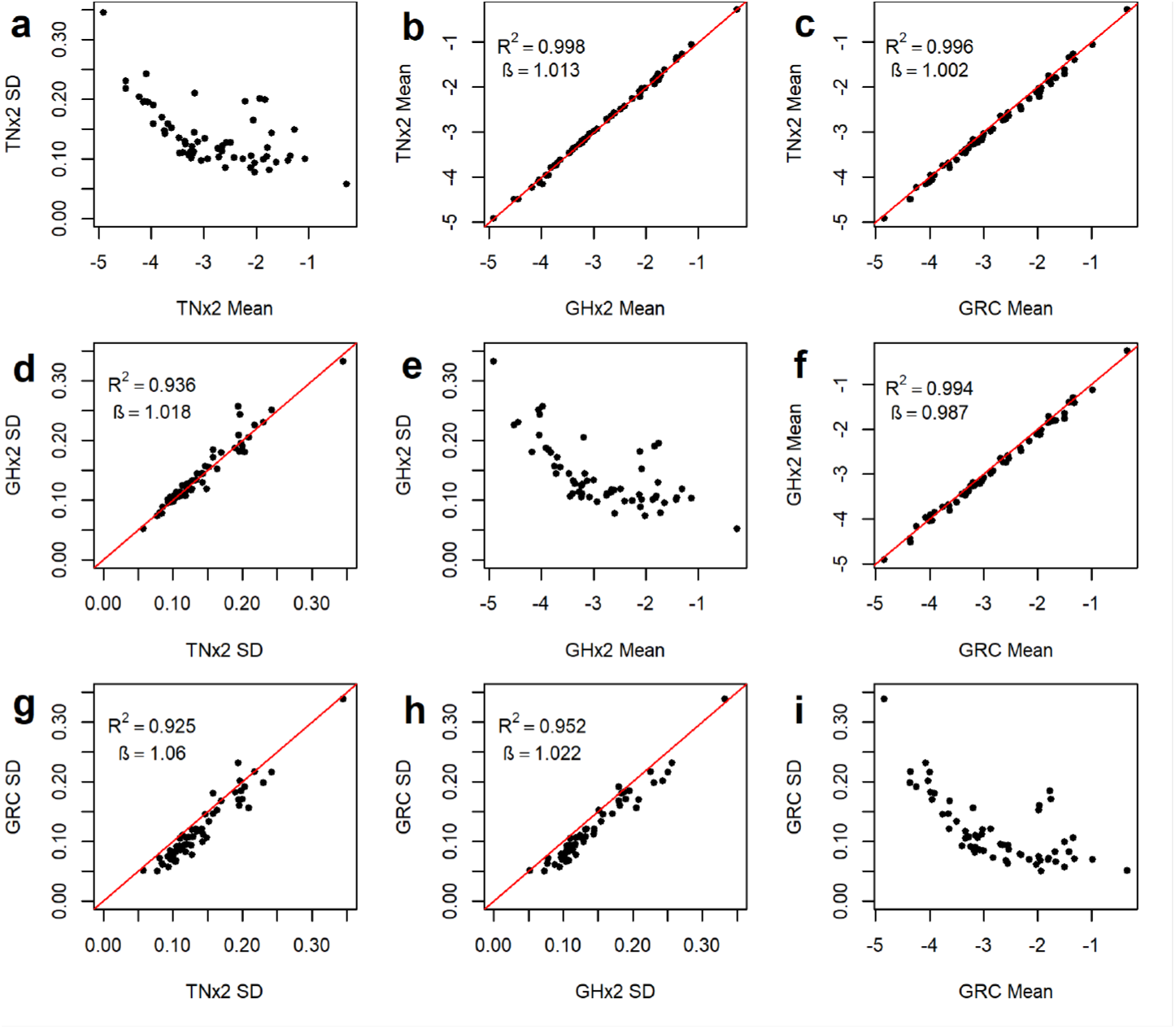
Comparison of means and standard deviations (SD) of log_10_ relative abundances using three sample preservation methods. Diagonal graphs (top left to bottom right) represent standard deviation plotted against the mean for TNx2 (**a**), GHx2 (**e**) and the GRC method (**i**). Above the diagonal are the plots of means against each other for the three different preservations methods (**b, c** and **f**). Below the diagonal are the plots of standard deviations against each other for the three different preservations methods (**d, g, h**). Red lines represent the diagonal line of identity.

Fig. 5 shows a matrix plot of the first three principal components (PCs) of a PCA analysis on the log_10_ relative abundances, with the plots in the upper panel colored based on preservation method and plots in the lower panel colored by sampling round. The first two PCs separated the GRC method from TNx2 and GHx2 (upper panel), while there was no separation between TNx2 and GHx2 from the first two PCs or separation between the methods based on PC3. The first two PCs also provide a high degree of separation between the two sampling rounds, whereas there was little separation between the rounds based on PC3. However, sampling rounds 1 separated from sampling round 2 in the direction of both PC1 and PC2 being positive, whereas separation of preservation methods (in direction of GRC) was along the positive PC1 and negative PC2 axes.

**Fig. 5.**
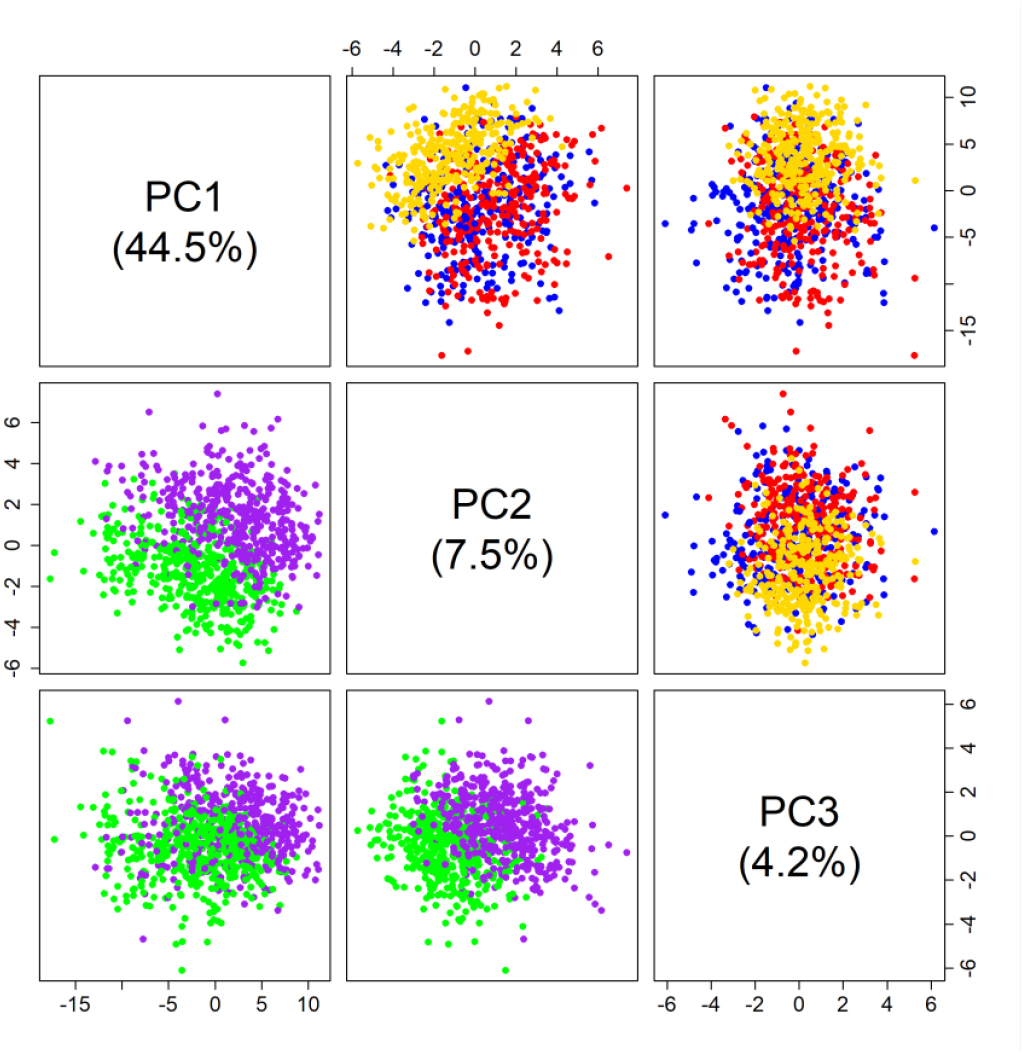
Matrix plot of first three PCs of log_10_ relative abundances. Points are colored by preservative method in the upper panel (gold is GRC, blue is TNx2 and red is GHx2) and sampling round in the lower panel (round 1 is green and round 2 is purple).

Fig. 6 plots the correlations of log_10_ relative abundances between samples for each microbial taxon, within a sampling round, for pairs of preservation methods. Results from the first round (Fig. 6 a,c,e) and the second round (Fig. 6 b,d,f) are plotted separately. The correlations of the taxa between GRC and TNx2 were similar to the correlations between GRC and GHx2 for both sampling rounds (Fig. 6 a,b), although the average correlation between GHx2 and GRC (0.69 ± 0.13) were slightly larger than between TNx2 and GRC (0.65 ± 0.13). For the other comparisons (GHx2 compared to GRC benchmarking against TNx2 (Fig. 6 c,d) and TNx2 compared to GRC benchmarking against GHx2 (Fig. 6 e,f)), the correlations were similar, suggesting a strong correspondence between the methods. The average correlation between TNx2 and GHx2 was 0.69 ± 0.12 and the overall average correlation was 0.68 ± 0.13. The correlations in log_10_ relative abundances across all comparisons was greater than 0.5 for most taxa, except for *Methanobrevibacter* and *Escherichia* which were the only genera below 0.5 in all cases. Correlations tended to be greater for gram-negative bacteria with the majority being between 0.7 and 0.95, whereas the correlations for gram-positive bacteria were smaller and mostly between 0.5 and 0.75. The ranking of the taxa in terms of their correlations was in general consistent between methods within a round (e.g., *Fibrobacter* was consistently the most correlated taxon between methods). However, there were a few exceptions with the correlations between TNx2 and GHx2 being much greater than the correlations between GRC and the two preservative solutions for *Methanobrevibacter* in sampling round 2 but much lower for *Lactococcus* in sampling round 1.

**Fig. 6.**
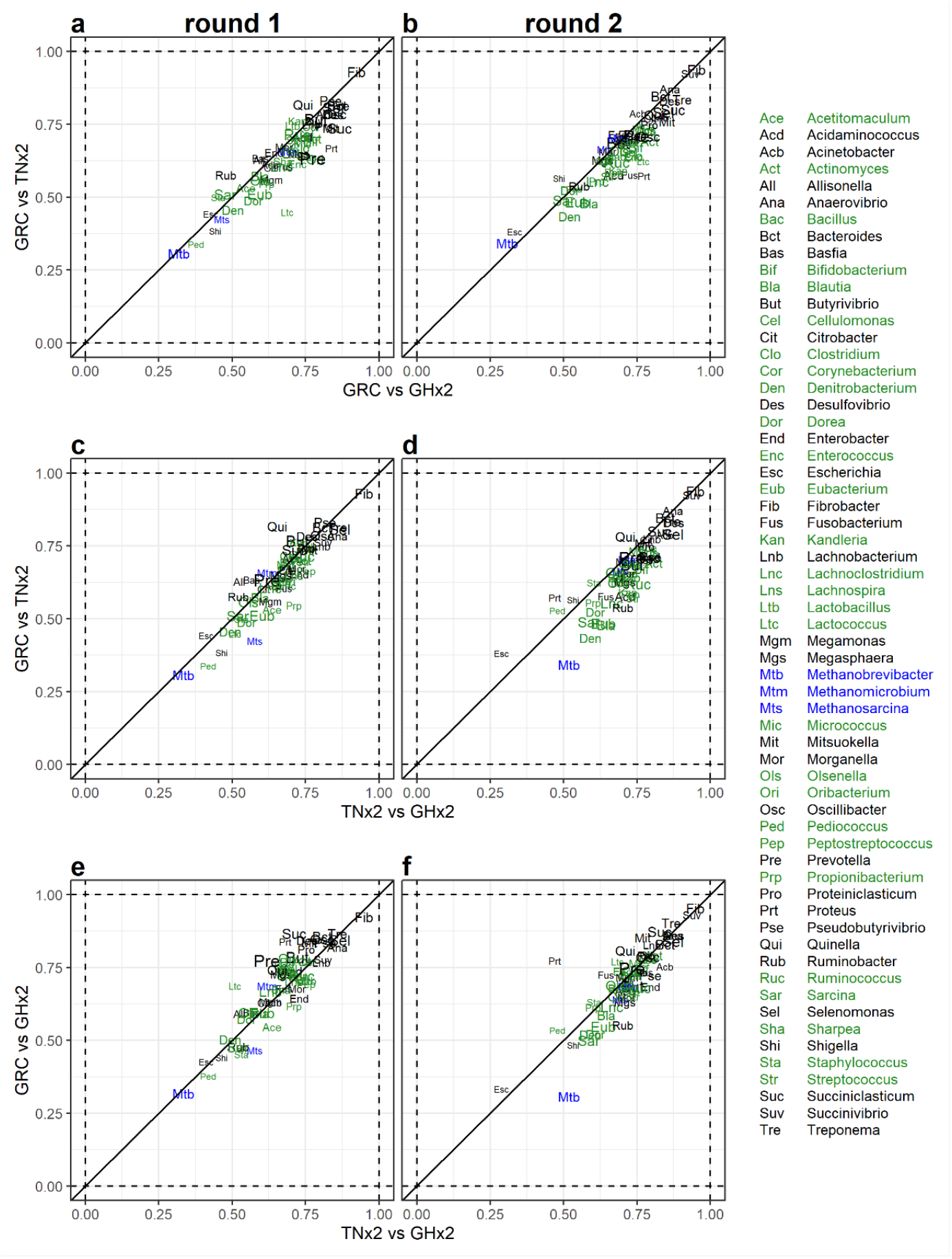
Pairwise correlation of log_10_ relative abundances for each microbial taxa between different preservation methods within sampling rounds. Correlations for sampling round 1 are given in the first column (**a**,**c**,**e**) and correlations for sampling round 2 are given in the second column (**b**,**d**,**f**). Each label in the scatter plots gives the pairwise correlation estimate (between the two methods on the *x*-axis compared to the two methods marked on the *y*-axis) in log_10_ relative abundances of each taxon for each sampling round. Taxon labels are colored based on whether the taxon is archaeal (blue), bacterial with a gram-positive wall (black) or bacterial with a gram-negative wall (green). The size of the taxon labels on the plots are proportional to the mean relative abundance of each taxon.

Fig. 7 plots, for each microbial taxon, the correlation of log_10_ relative abundances between samples collected on the same animal at round 1 and round 2 and preserved using either TNx2 or GHx2 compared to GRC. The correlations between sampling rounds were, in general, between 0 and 0.5 for all methods and most taxa, with only two genera (*Ruminobacter* and *Streptococcus*) having small negative correlations in same cases. The average correlation across taxa was 0.15 ± 0.09 for TNx2, 0.19 ± 0.10 for GHx2, 0.18 ± 0.12 for GRC, and 0.17 ± 0.11 overall. There were no patterns whereby more common genera (or gram-negative bacteria) were closer to the diagonal line and rarer genera (or gram-positive bacteria) were further from the diagonal line. However, the three archaeal taxa tended to fall below the diagonal line indicating greater correlations between rounds for the GRC method compared to TNx2 and GHx2.

**Fig. 7.**
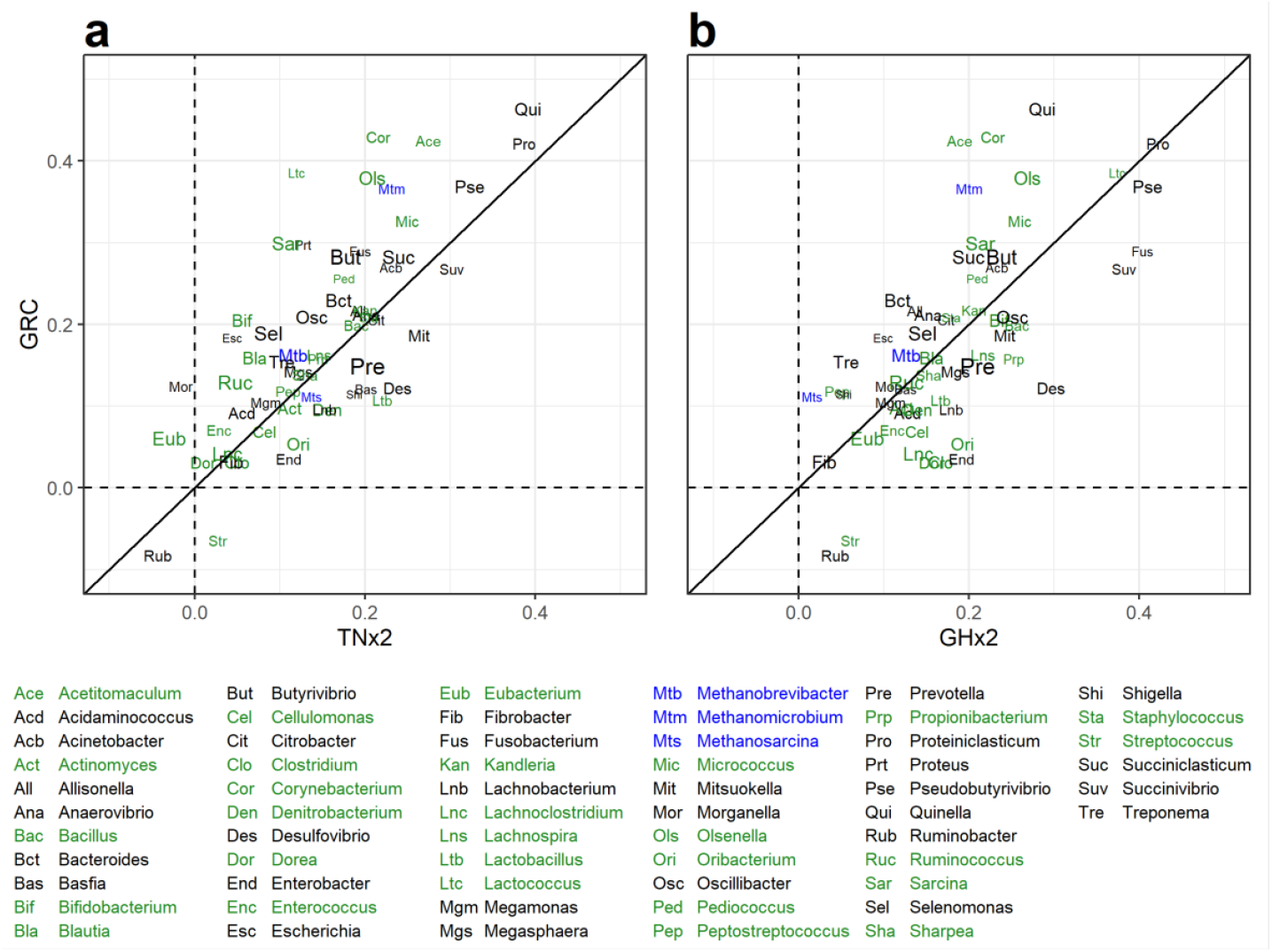
Correlation of log_10_ relative abundance between sampling round 1 and 2 for samples preserved using different methods. **a** Correlation estimates between rounds for samples preserved using GRC compared to samples preserved using TNx2. **b** Correlation estimates between rounds for samples preserved using GRC compared to samples preserved using GHx2. Taxon labels are colored according to whether they are archaeal (blue), bacterial with a gram-positive wall (black) or bacterial with a gram-negative wall (green). The size of the taxon labels on the plots are proportional to the mean relative abundance of each taxon.

## Discussion

The objective of this research was to develop a low-cost preservation method that enables access to high-throughput DNA extraction from rumen samples. We used three different preservative solutions for rumen samples collected from sheep, then generated RMC profiles using RE-RRS, benchmarking against the GRC method. Our results showed that using pure ethanol to preserve rumen samples resulted in poor DNA quantity and quality, and large differences in the RMC profile compared to the GRC method, which has been successfully used in a global project to survey RMCs [13]. Therefore ethanol at the final concentration used in this study is not recommended for preserving rumen samples for subsequent microbiome analyses. In contrast, the two preservative solutions that we proposed, TNx2 and GHx2, resulted in DNA of high quality and integrity, and RMC profiles with a high degree of similarity to those obtained using the GRC method.

### Rumen content and DNA integrity

Variations in sampling methods, sample processing and storage, and DNA extraction protocols used in this trial impacted rumen phase content and DNA integrity. For the GRC method, rumen samples were taken directly from the rumen via stomach tubing or at slaughter, and the freeze-dried sample was used for DNA extraction, which makes the solid phase more represented compared to alternative preservation methods. For the three preservative solutions, the rumen sample was subsampled twice using a pipette (firstly from the 30 mL pottle of rumen contents, and then later from the 8 mL vial for DNA extraction). This method results in the sample consisting primarily of the liquid phase of the rumen. This may explain the larger DNA yield observed in the GRC method, where the predominant bacterial mass is adhered to the ruminal fibre [34].

Besides the rumen sample phase effect on DNA, it is known that the DNA extraction method used may generate different DNA yield, quality, and integrity [16]. The GRC method uses the PCQI protocol, which has previously been tested in sheep rumen samples with a DNA yield of 824 µg/g per DM being obtained [16], which is smaller than the average DNA yield observed in this study (988 µg/g of DM). DNA extractions for the preservative solutions followed the PCQI protocol with some modifications since these solutions predominantly have a conservation and lysis function. Of the three solutions, GHx2 resulted in the greatest DNA yield (857 µg/g of DM), which was slightly higher than the DNA yield observed in Henderson et al. [16], while TNx2had a lower DNA yield (511 µg/g of DM). This suggests the solutions, in addition to feed type, time off feed and the rumen fill, had varying impacts on DNA preserving and extraction and consequently on its yield. Studies have suggested that 6 M GuHCl/EDTA solution is able to preserve bacterial genetic materials as well as freezing, providing additional conveniences [35]. Methodologically, TNx2 and GHx2 were more comparable between each other, and handled similarly compared to EtOH (**Table *1*** Table 2). The major differences in GHx2 from TNx2 were the replacement of a) the NaCl salt by GuHCl and b) the surfactant SDS with Tween 20 and Triton x100.

The way in which samples were processed prior to DNA extraction (e.g., grinding) and the storage efficiency of each method in preserving samples over time are secondary factors that may also have led to differences in our results [17]. Freezing and freeze drying are widely used as a standard method for long-term storage of different organisms [36]. However, it is known that freezing can result in injury or cellular damage and changes in cell membrane permeability, while changes have been observed in frozen and freeze-dried samples (e.g., DNA yield, bacterial abundances) compared with fresh samples [12, 37-39].

### Sequencing and RMC profiles

Hess et al. [10] developed the RE-RRS approach using *Pst*I to study RMC profiles using the GRC method. These authors observed more reads per sample (2.7 million) compared to the GRC samples in this trial (1 million), which is due to Hess et al. [10] using fewer samples per lane (118) compared to this study (302). The authors suggested through a sensitivity analysis that up to 2000 samples could theoretically be sequenced in a single lane on an Illumina HiSeq 2500 to obtain realiable RMC information, and the 302 samples per lane used in this study is well within that threshold.

Our sequencing results revealed that RE-RRS with *Pst*I restriction enzyme was suitable for samples preserved using our proposed preservative solutions. Compared to the GRC method, the number of reads generated from sequencing was similar for TNx2 and slightly lower for GHx2. However, a larger proportion of these reads were assigned a genus-level taxonomy and the correlations for individual taxa in the resulting RMC profiles were similar between GRC and the two lysis buffer-based solutions. These differences are likely to be due to differential DNA extraction from the taxa present and the mix of genomes in the reference database used for analysis. EtOH proved to be an anomaly, with significantly fewer reads and more failed samples. This may have been due to poor sample quality, as using DNA with some level of degradation for in library preparation reduces the efficiency of sequencing and generates unrepresentative data on the HiSeq 2500 platform.

The relative abundance of genera was found to differ between preservation methods. The five gram-negative bacteria *Anaerovibrio, Bacteroides, Prevotella, Selenomonas* and *Succiniclasticum* were in lower abundance, and gram-positive bacteria had similar or larger abundances in GRC preserved samples compared to TNx2 and GHx2. These changes in relative abundances may relate to the different characteristics of the cell wall between gram-positive and gram-negative bacteria. Gram-positive bacteria have thicker walls and these are likely to be more resistant to breakage than those of gram-negative bacteria. Gram-negative bacteria could be biased against if they lysed during storage and their DNA degraded. DNA extraction from any bacterium, gram-positive or gram-negative, could be biased for or against if the preservative decreased or increased, respectively, the structural integrity of the wall. However, because the final data are proportional, the contribution of different biases are not able to be elucidated. The least abundant genus, *Shigella*, was the only taxon not affected by preservation method, which is likely due to this microbe representing contamination as it is not naturally present in the rumen [4].

The RMC profiles of samples preserved with TNx2 and GHx2 were very similar, with comparable mean relative abundances across the majority of taxa and little separation in the PCA analysis. Despite differences in mean relative abundances between the TNx2 and GHx2 samples with the GRC samples, the correlations in log_10_ relative abundances of individual genus-level taxa between these three methods were similar (Fig. 6). This suggests that the ranking of animals within a method is consistent even if there are shifts in mean abundances, which has some important practical implications. One is that using TNx2 or GHx2 instead of the GRC method should have minimal impact on trait prediction in breeding applications. This scenario, nevertheless, assumes that samples have been preserved using a single method. If this is not the case, some level of normalization will be required, such as the cohort standardization of the RMC profile used in Hess et al. [10], to enable robust predictions across preservation methods.

The sampling round also impacted RMC profiles for all methods, with significant changes in mean relative abundance across a mixture of gram-positive, gram-negative and archaeal taxa. The correlations between individual taxon abundances were lower between rounds for the same method (Fig. 7) compared to different methods within a round (Fig. 6), which highlights that time of sampling for this study has a larger impact on RMC profiles than the preservation method. Differences in RMC profiles between rounds are possibly driven by differences in the feed, which are suspected to be major drivers of RMC variation over time in sheep grazing pasture [40]. Nevertheless, the correlations between rounds within a method (Fig. 7) were similar across the three preservation methods (TNx2, GHx2 and GRC method), which further suggests a high degree of concordance between these preservative solutions and the GRC method.

### Comparison of preservative solutions with GRC method

The preservative solutions we tested provide several practical advantages over the GRC method. The first is that samples can be kept chilled (fridge temperature) for a few days after collection before freezing, whereas samples are frozen at collection under the GRC method, which requires a freezer or cold substances (e.g., liquid nitrogen, dry ice) on hand for immediate freezing. This adds additional cost and handling to field and on-farm sample collections. The main advantage is that the preservative solution protocol bypasses the pre-processing steps of freeze drying and grinding used in the GRC method. These pre-processing steps are a bottleneck in the sample processing as they require laboratory staff with technical expertise, take ∼12 days to process a batch of 186 samples (∼8 days freeze drying and ∼1.5 days grinding), and a limited number of samples can be processed at a given time, depending on the quantity of freeze drying and grinding machines available. Using the preservative solutions would therefore accelerate the DNA extraction process and, coupled with RE-RRS, enable high-throughput generation of RMC profiles.

Throughout this study, the GRC method has assumed the role as the “gold standard”, and the performance of the preservative solutions tested in this study were primarily evaluated based on their similarity to the GRC method. In this evaluation, we are testing the preservative properties of the methods and not the ability to extract DNA since the DNA extraction protocol was similar between all methods. Our results indicate that for TNx2 and GHx2, there was little bias in the mean log_10_ relative abundances when averaging across all taxa, but more variation compared to the GRC method. This suggests that TNx2 and GHx2 are both viable alternatives to the GRC method for preserving rumen samples, but more samples need to be collected to obtain the same power to correlate the rumen microbiome with traits. The GRC method, however, does not represent a true “gold standard”, as it is unknown how well it captures the true RMC, and previous studies have found differences between frozen and fresh samples in several organisms [12, 37]. The impact of using the proposed alternative preservation methods should therefore be evaluated in other applications of RMC profiles (e.g., for trait prediction), which is the focus of current research.

Several important factors need to be considered when selecting one of the TNx2 and GHx2 preservative solutions as an alternative to the GRC method. Firstly, our results suggest that GHx2 yielded greater quantities of DNA and resulted in RMC profiles that were more similar to the GRC method compared to TNx2, evidenced by slightly higher correlations in log_10_ relative abundances of individual taxa with the GRC method (Fig. 6). This implies that the GHx2 solution provides more comparable results to the GRC method than TNx2. However, an important practical consideration is the ease of obtaining raw chemicals used in the solutions and their inherent toxicity. Toxic chemicals require greater safety precautions, additional operator training, and can complicate the transportation of samples, both domestically and internationally. In comparing TNx2 with GHx2, the key component is the salt used in the lysis solution. GHx2 uses GuHCl as its salt base, which is not only more expensive and difficult to source but is also more toxic than NaCl used in TNx2. In our view, these practical advantages of TNx2 outweigh the marginal improvement in the RMC profile obtained using GHx2. We therefore recommend the TNx2 solution as an alternative to the GRC method for industry application.

## Conclusions

We explored the use of three solutions as alternative preservation methods to the Global Rumen Census method that involves costly freeze drying and grinding steps. Of the three solutions, TNx2 and GHx2 were found to be viable alternatives as they provided high quality DNA and showed relatively little bias in the relative abundance of the microbes compared to the GRC method, although larger variances in relative abundances were observed in samples from grazing sheep. However, the variance due to different time-points was greater than that attributed to preservation methods. This suggests, given appropriate cohort adjustment, that RMC profiles from different preservation methods could be combined for breeding purposes. The use of these preservative solutions significantly reduces both labor and costs associated with processing rumen samples for microbial sequencing and, if used in conjunction with RE-RRS, would enable low-cost and high-throughput generation of RMC profiles. This could greatly enhance the implementation of RMC as a tool for breeding in the global livestock industry.

## Abbreviations

DM: Dry matter
GRC: Global Rumen Census
GuHCl: Guanidine hydrochloride
PCA: Principal component analysis
PC: Principal component
RB: Reference-based
RE-RRS: Restriction enzyme-reduced representation sequencing
RMC: Rumen microbial community
SDS: Sodium dodecyl sulfate

## Additional Files

**Additional file 1. Supplementary tables and figures**. Table S1) Dry matter (DM) of the GRC method samples using the freeze dryer. Table S2) DNA yield and DNA quality for rumen samples preserved using the GRC method, TNx2, GHx2 and EtOH. Table S3) DNA yield and quality of sheep rumen samples using four sample preservation methods (expanded). Table S4) Differences in relative abundance of genera using four different preservation methods. Fig. S1) PCR plate layout for the four different libraries.

**Additional file 2. Supplementary data**. Data S1) Sample and library information of 1208 samples preserved using the four preservation methods and 32 positive controls sequenced using RE-RRS. Data S2) Number of reads generated from RE-RRS for 1208 samples preserved using the four preservation methods and 72 control sample (32 positive and 40 negative). Data S3) Count matrix of reads assigned to genus level taxonomy using the RB approach for 1208 samples preserved using the four preservation methods and 72 control sample (32 positive and 40 negative).

**Additional file 3. Supplementary figures for visual assessment of sheep rumen samples using different preservation methods**. Fig. S2) 8 mL vial containing the sheep rumen sample preserved with TNx2, GHx2 or EtOH. Fig. S3) 2 mL vials containing TNx2, GHx2 or EtOH and sheep rumen sample after the “bead-beating” step. Fig. S4) 2 mL vials containing sheep rumen samples preserved using TNx2, GHx2, EtOH and the GRC method after centrifugation. Fig. S5) Principal component analysis (PCA) of the log_10_ relative abundance matrix using the RB approach for all non-failed samples (> 100,000 reads) including positive control samples.

**Additional file 4. Supplementary Code**. R code used to produce the analysis.

## Acknowledgements

We acknowledge Brooke Bryson, Anna Boyd, Larissa Zetouni and Erin Waller for assistance with the collection and processing of the samples.

## Authors Contributions

JM and SR produced and reared the research flock and conceived the study while JM designed the trial. HH handled the samples in the lab for sequencing. JB, MH and TB jointly conducted the statistical analyses and drafted the manuscript. KD provided statistical support for the analyses. All authors helped interpret the results and revised the manuscript.

## Availability of Data and Material

Raw sequence files are available from the NCBI SRA database (BioProject ID: PRJNA791831). A temporary reviewer link is https://dataview.ncbi.nlm.nih.gov/object/PRJNA791831?reviewer=ulqggsue533i6ig4peecrnf69k. Lab data, metadata and processed RE-RRS data (read numbers and RB count matrix) are give in Additional file 2 and the R code used for the analysis is in Additional file 4.

## Declarations

### Ethics Approval

The use of experimental animals and protocols applied in this experiment were approved by the AgResearch Invermay (Mosgiel, NZ) Animal Ethics committee (approval number 14370).

### Consent for Publication

Not applicable

## References

1. Moraïs S, Mizrahi I: The road not taken: the rumen microbiome, functional groups, and community states. Trends in Microbiology 2019, 27:538–549.

2. Huws SA, Creevey CJ, Oyama LB, Mizrahi I, Denman SE, Popova M, Muñoz-Tamayo R, Forano E, Waters SM, Hess M: Addressing global ruminant agricultural challenges through understanding the rumen microbiome: past, present, and future. Frontiers in Microbiology 2018, 9:2161.

3. Goodacre R: Metabolomics of a superorganism. The Journal of Nutrition 2007, 137:259S–266S.

4. Seshadri R, Leahy SC, Attwood GT, Teh KH, Lambie SC, Cookson AL, Eloe-Fadrosh EA, Pavlopoulos GA, Hadjithomas M, Varghese NJ: Cultivation and sequencing of rumen microbiome members from the Hungate1000 Collection. Nature Biotechnology 2018, 36:359–367.

5. Morgavi D, Kelly W, Janssen P, Attwood G: Rumen microbial (meta) genomics and its application to ruminant production. Animal 2013, 7:184–201.

6. Ross EM, Moate PJ, Bath CR, Davidson SE, Sawbridge TI, Guthridge KM, Cocks BG, Hayes BJ: High throughput whole rumen metagenome profiling using untargeted massively parallel sequencing. BMC Genetics 2012, 13:53.

7. Delgado B, Bach A, Guasch I, González C, Elcoso G, Pryce JE, Gonzalez-Recio O: Whole rumen metagenome sequencing allows classifying and predicting feed efficiency and intake levels in cattle. Scientific Reports 2019, 9:11.

8. Mizrahi I, Jami E: The compositional variation of the rumen microbiome and its effect on host performance and methane emission. Animal 2018, 12:s220–s232.

9. Tapio I, Snelling TJ, Strozzi F, Wallace RJ: The ruminal microbiome associated with methane emissions from ruminant livestock. Journal of Animal Science and Biotechnology 2017, 8:7.

10. Hess MK, Rowe SJ, Van Stijn TC, Henry HM, Hickey SM, Brauning R, McCulloch AF, Hess AS, Kirk MR, Kumar S, et al: A restriction enzyme reduced representation sequencing approach for low-cost, high-throughput metagenome profiling. PLoS One 2020, 15:e0219882.

11. Elshire RJ, Glaubitz JC, Sun Q, Poland JA, Kawamoto K, Buckler ES, Mitchell SE: A robust, simple genotyping-by-sequencing (GBS) approach for high diversity species. PLoS One 2011, 6:e19379.

12. Metzler-Zebeli BU, Lawlor PG, Magowan E, Zebeli Q: Effect of freezing conditions on fecal bacterial composition in pigs. Animals 2016, 6:18.

13. Henderson G, Cox F, Ganesh S, Jonker A, Young W, Collaborators GRC, Janssen PH: Rumen microbial community composition varies with diet and host, but a core microbiome is found across a wide geographical range. Scientific Reports 2015, 5:14567.

14. Kittelmann S, Pinares-Patiño CS, Seedorf H, Kirk MR, Ganesh S, McEwan JC, Janssen PH: Two different bacterial community types are linked with the low-methane emission trait in sheep. PLoS One 2014, 9:e103171.

15. Rius AG, Kittelmann S, Macdonald KA, Waghorn GC, Janssen PH, Sikkema E: Nitrogen metabolism and rumen microbial enumeration in lactating cows with divergent residual feed intake fed high-digestibility pasture. Journal of Dairy Science 2012, 95:5024–5034.

16. Henderson G, Cox F, Kittelmann S, Miri VH, Zethof M, Noel SJ, Waghorn GC, Janssen PH: Effect of DNA extraction methods and sampling techniques on the apparent structure of cow and sheep rumen microbial communities. PLoS One 2013, 8:e74787.

17. Granja-Salcedo YT, Ramirez-Uscategui RA, Machado EG, Duarte Messana J, Takeshi Kishi L, Lino Dias AV, Berchielli TT: Studies on bacterial community composition are affected by the time and storage method of the rumen content. PLoS One 2017, 12:e0176701.

18. Hammond PM: Protein purification. In Encyclopedia of microbiology. Volume 3. Edited by Lederberg J. San Diego: Academic Press Inc; 1992: 451–460

19. Silhavy TJ, Kahne D, Walker S: The bacterial cell envelope. Cold Spring Harbor Perspectives in Biology 2010, 2:a000414.

20. Damberg P, Jarvet J, Gräslund A: Micellar Systems as Solvents in Peptide and Protein Structure Determination. In Methods in Enzymology. Volume 339. Edited by James TL, Dötsch V, Schmitz U: Academic Press; 2001: 271–285

21. Pramanick D, Forstova J, Pivec L: 4 M guanidine hydrochloride applied to the isolation of DNA from different sources. FEBS letters 1976, 62:81–84.

22. Shokralla S, Singer GA, Hajibabaei M: Direct PCR amplification and sequencing of specimens’ DNA from preservative ethanol. Biotechniques 2010, 48:305–306.

23. Montgomery GW, Sise JA: Extraction of DNA from sheep white blood cells. New Zealand Journal of Agricultural Research 1990, 33:437–441.

24. Rowe SJ, Hickey SM, Jonker A, Hess MK, Janssen P, Johnson T, Bryson B, Knowler K, Pinares-Patiño C, Bain W, et al: Selection for divergent methane yield in New Zealand sheep - a ten-year perspective. Proceedings of the Association for the Advancement of Animal Breeding and Genetics 2019, 23:306–309.

25. Pinares-Patiño CS, Hickey SM, Young EA, Dodds KG, MacLean S, Molano G, Sandoval E, Kjestrup H, Harland R, Hunt C, et al: Heritability estimates of methane emissions from sheep. Animal 2013, 7:316–321.

26. Jonker A, Hickey SM, Rowe SJ, Janssen PH, Shackell GH, Elmes S, Bain WE, Wing J, Greer GJ, Bryson B, et al: Genetic parameters of methane emissions determined using portable accumulation chambers in lambs and ewes grazing pasture and genetic correlations with emissions determined in respiration chambers1. Journal of Animal Science 2018, 96:3031–3042.

27. Herten K, Hestand MS, Vermeesch JR, Van Houdt JK: GBSX: a toolkit for experimental design and demultiplexing genotyping by sequencing experiments. BMC Bioinformatics 2015, 16:73.

28. Martin M: Cutadapt removes adapter sequences from high-throughput sequencing reads. EMBnet Journal 2011, 17:10–12.

29. Kumar S: Physiology of rumen bacteria associated with low methane emitting sheep. Doctoral. Massey University, 2017.

30. R Core Team: R: A Language and Environment for Statistical Computing. Vienna, Austria: R Foundation for Statistical Computing; 2020.

31. Bates D, Maechler M, Bolker B, Walker S: Fitting linear mixed-effects models using lme4. Journal of Statistical Software 2015, 67:1–48.

32. Luo D, Ganesh S, Koolaard J: predictmeans: Calculate predicted means for linear models (R package version 1.0.6). 2021.

33. Antunes LCS, Poppleton D, Klingl A, Criscuolo A, Dupuy B, Brochier-Armanet C, Beloin C, Gribaldo S: Phylogenomic analysis supports the ancestral presence of LPS-outer membranes in the Firmicutes. eLife 2016, 5:e14589.

34. Vaidya JD, van den Bogert B, Edwards JE, Boekhorst J, Van Gastelen S, Saccenti E, Plugge CM, Smidt H: The effect of DNA extraction methods on observed microbial communities from fibrous and liquid rumen fractions of dairy cows. Frontiers in Microbiology 2018, 9:92.

35. Ribeiro RM, Souza-Basqueira Md, Oliveira LCd, Salles FC, Pereira NB, Sabino EC: An alternative storage method for characterization of the intestinal microbiota through next generation sequencing. Revista do Instituto de Medicina Tropical de São Paulo 2018, 60.

36. Perry SF: Freeze-drying and cryopreservation of bacteria. Molecular Biotechnology 1998, 9:59–64.

37. Hsu JT, Fahey GC, Jr.: Effects of Centrifugation Speed and Freezing on Composition of Ruminal Bacterial Samples Collected from Defaunated Sheep. Journal of Dairy Science 1990, 73:149–152.

38. Nei T: Freezing and freeze-drying of microorganisms. Cryobiology 1964, 1:87–93.

39. Spanghero M, Chiaravalli M, Colombini S, Fabro C, Froldi F, Mason F, Moschini M, Sarnataro C, Schiavon S, Tagliapietra F: Rumen inoculum collected from cows at slaughter or from a continuous fermenter and preserved in warm, refrigerated, chilled or freeze-dried environments for in vitro tests. Animals 2019, 9:815.

40. Moon CD, Carvalho L, Kirk MR, McCulloch AF, Kittelmann S, Young W, Janssen PH, Leathwick DM: Effects of long-acting, broad spectra anthelmintic treatments on the rumen microbial community compositions of grazing sheep. Scientific Reports 2021, 11:3836.

