## Additional file 3 for "Low-cost sample preservation methods for high-throughput processing of rumen microbiomes"

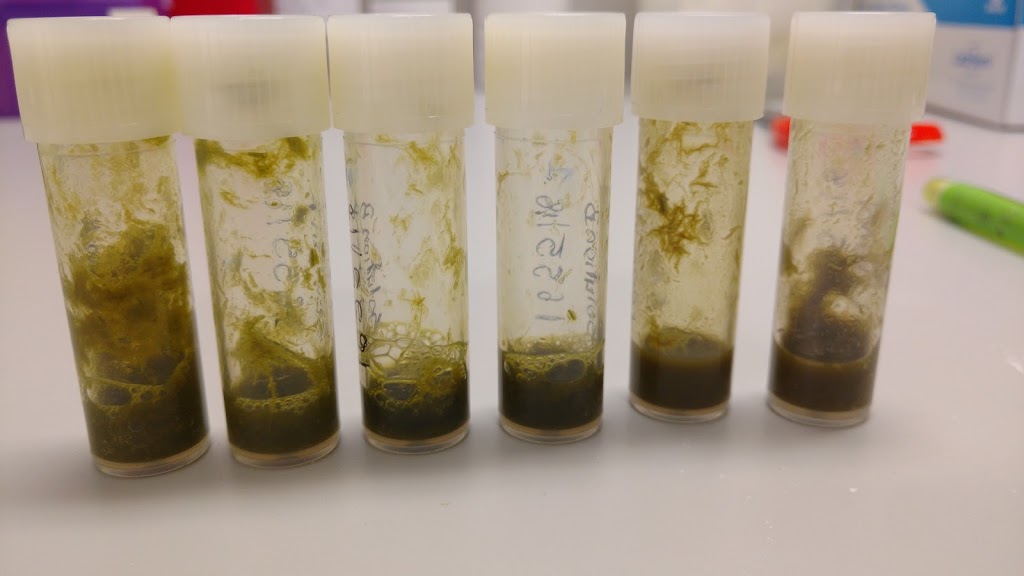

**(a)**

**(b)**

**(c)**

**(b)**

**(c)**

**(a)**
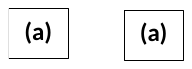

**Fig. S2** 8 mL vials containing the same sheep rumen sample preserved with (a) TNx2, (b) GHx2 or (c) EtOH.

**(a)**
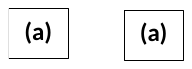

**(c)**

**(b)**

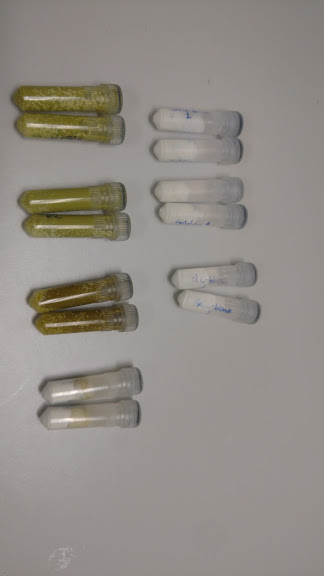

**Fig. S3** 2 mL vials containing (a) TNx2, (b) GHx2, or (c) EtOH with the same sheep rumen sample after the “bead-beating” step.

| 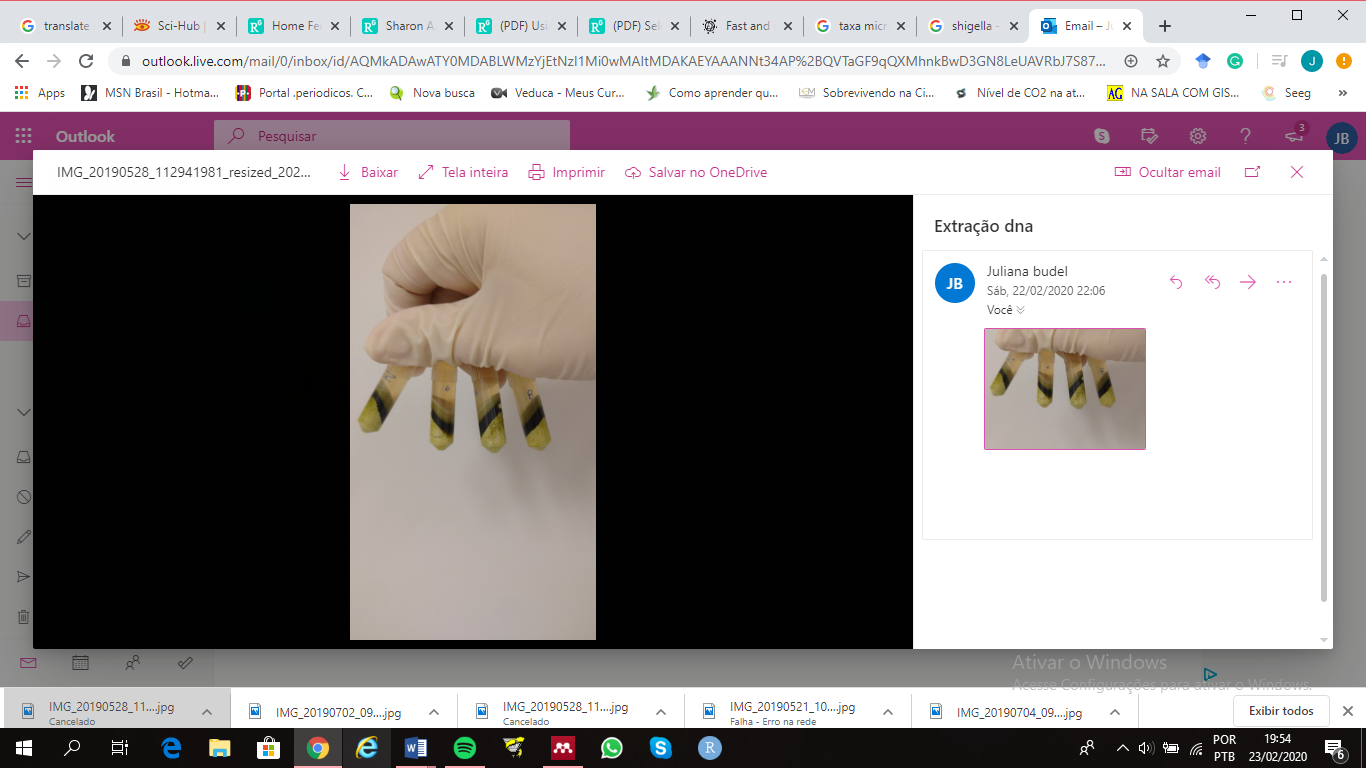 | 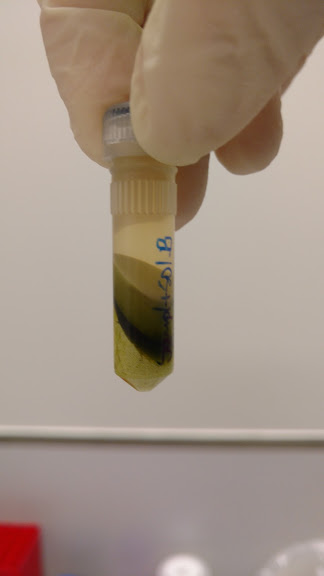 | 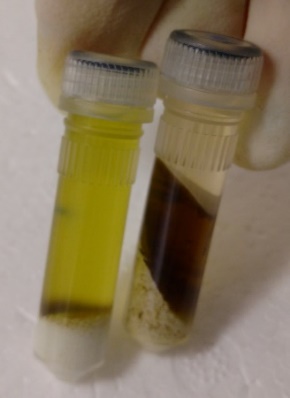 | 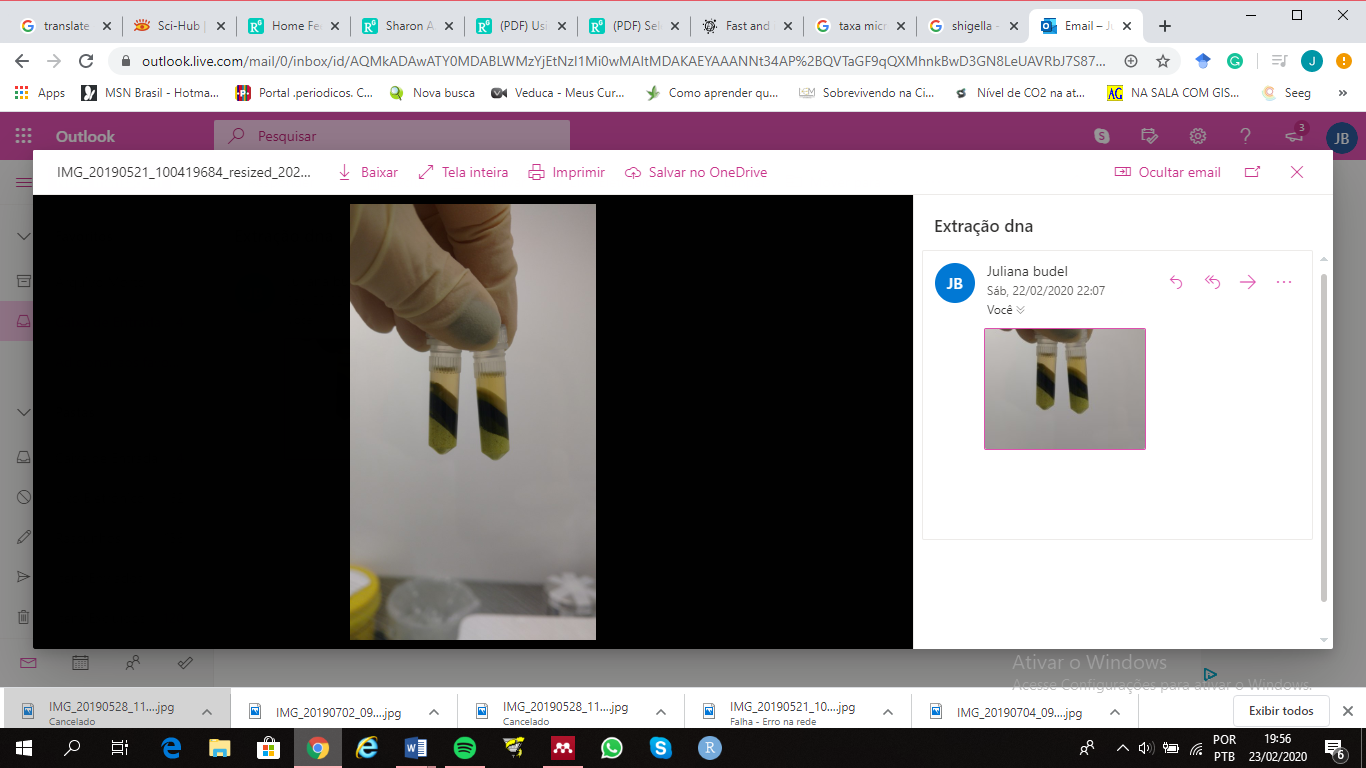 |
| --- | --- | --- | --- |
| **(a)** | **(b)** | **(c)** | **(d)** |

**Fig. S4** 2 mL vials containing the same sheep rumen samples preserved using (a) TNx2, (b) GHx2, (c) EtOH, and (d) the GRC method after centrifugation.

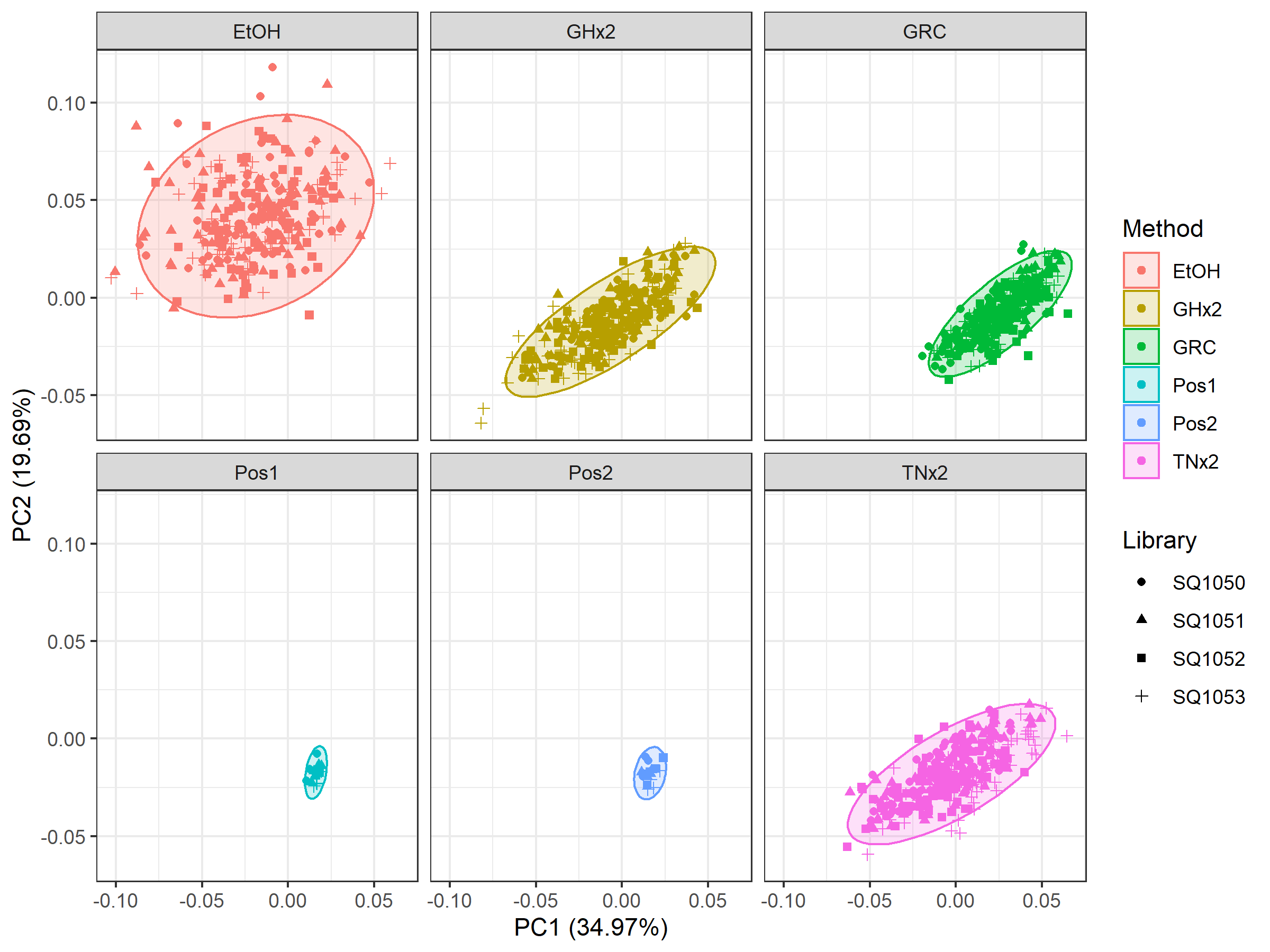

**Fig. S5** Principal component analysis (PCA) of the log_10_ relative abundance matrix using the RB approach for all non-failed samples (>100k reads) including positive control samples. The 32 positive controls samples were split equally across two individuals (labelled Pos1 and Pos2) and were preserved using the GRC method.
